# Efficacy of Centers of Biomedical Research Excellence (CoBRE) Grants to Build Research Capacity in Underrepresented States

**DOI:** 10.1101/2023.08.02.551624

**Authors:** Michael D. Schaller

## Abstract

Federal funding for research has immediate and long-term economic impact. Since federal research funding is regionally concentrated and not geographically distributed, the benefits are not fully realized in some regions of the country. The Established (previously Experimental) Program to Stimulate Competitive Research (EPSCoR) programs at several agencies, e.g. the National Science Foundation, and the Institutional Development Award (IDeA) program at the National Institutes of Health were created to increase competitiveness for funding in states with historically low levels of federal funding. The Centers of Biomedical Research Excellence (CoBRE) award program is a component of the IDeA program. The CoBRE grants support research core facilities to develop research infrastructure. These grants also support the research projects of junior investigators, under the guidance of mentoring teams of senior investigators, to develop human resources at these institutions. Few studies have assessed the effectiveness of these programs. This study examines the investment and outcomes of the CoBRE grants from 2000 through 2022. The maturation of junior investigators into independently funded principal investigators is comparable to other mentoring programs supported by NIH. The investment in research cores resulted in substantial research productivity, measured by publications. Despite the successes of individual investigators and increase research infrastructure and productivity, the geographic distribution of federal and NIH research dollars has not changed. These results will be informative in consideration of policies designed to enhance the geographic distribution of federal research dollars.

## Introduction

The modern US research and development system arose after World War II. Inspired by the wartime scientific breakthroughs fueled by government investment, the idea that science is vital for human health and the economy took hold (1). Two novel components of the emerging policy were the funding of basic research in academia by the federal government and the provision of federal scholarships and fellowships to support training in science (2, 3). Implementation of this policy led to the development of the grant funding system at the National Institutes for Health (NIH) and creation of the National Science Foundation (NSF). The new laws codifying the policies directed building scientific capacity in all geographic areas and implementation fell to the agency administrations (3). Initially, funding was strongly skewed to a small number of institutions. The distribution of federal funding broadened over time, but is still regionally concentrated, which can be an impediment to economic development and training the scientific workforce in some areas of the country (2).

Investment in research and development has an economic impact in multiple ways. In the immediate term, research dollars support salaries of researchers, who buy goods and services that support the local economy. In 2008, $1 of research funding from the NIH produced an average of a $2.21 increase in the state economy (4). In the longer term, research provides economic development. Innovation is critical for creating sustained, long-term economic growth in modern economies and public funding of research drives innovation (5, 6). The results of federally funded research and development at academic institutions spills over to businesses in several ways (6). Knowledge transfer is the obvious spillover and can be documented by tracing citations in patents. NIH-funded research leads to patents, with 8.4% of grants directly leading to patents and 31% of grants indirectly leading to patents by producing papers that contribute to the intellectual basis of patents, as evidenced by citations (7). Public investment in research leads to additional investment from private sources, although there is a lag of 5 to 10 years (8–10). The return on investment of publicly funded research in terms of the output of bioscience industries is estimated from $1.70 to $3.15 per every $1 spent by NIH (9). In addition to the generation of knowledge, academic research provides the training ground for the next generation of scientists, i.e. the future workforce in industry (6). The movement of newly trained researchers from academia into industry develops networks that are important for optimization of economic development (6, 11). For this and other reasons, there is a geographic limitation on the diffusion of knowledge from academic institutions, particularly public institutions, and industry (12). The historical regional concentration of federally supported research and development provides an economic advantage in some regions, while other regions lack this opportunity (2, 13).

Congressional mandates led federal funding agencies to create programs to build research capacity, both research infrastructure and human resources, in regions that are typically underfunded to redress the problem of inequitable distribution of federal research dollars. In 1978, the NSF created the Experimental Program to Stimulate Competitive Research (EPSCoR) to meet this challenge. Other federal agencies have also created EPSCoR like programs. In 1993, NIH established the Institutional Development Award (IDeA) program to address the same issue in NIH funding. This is particularly important since ∼56% of federal research expenditures in institutions of higher education come from the Department of Health and Human Services (HHS), which administers NIH. The IDeA program has three main mechanisms to build research infrastructure: the IDeA Networks of Biomedical Research Excellence (INBRE) program, the Centers of Biomedical Research Excellence (CoBRE) program (initiated in 2000) and the IDeA Networks for Clinical and Translational Research (IDeA-CTR) program (initiated in 2011). The INBRE program supports statewide networks for the support of faculty research, student engagement in research and research infrastructure. The CoBRE program builds capacity in an area of research by supporting research projects by junior faculty, mentoring to facilitate faculty success and improvement of research infrastructure in the scientific area. The IDeA-CTR grants aim to support the development of infrastructure to perform clinical and translational research, increase competitiveness for clinical and translational research programs and establish collaborative efforts across IDeA institutions to better serve their populations.

Few studies have addressed the effectiveness of these congressionally mandated efforts to build research infrastructure and to balance the distribution of the federal investment in research and development. This study addresses the NIH CoBRE program and measures the investments made and the scientific outcomes of the program. The program succeeds in producing the primary desired outcomes of mentoring junior faculty to independence and providing infrastructure support for research. Despite these successes, there is little evidence for a sustained impact in changing the geographical distribution of federal funding.

## Methods

### Identification of CoBRE Awards

Institutions eligible for CoBRE grants are in IDeA states, which include Alaska, Arkansas, Delaware, Hawaii, Idaho, Kansas, Kentucky, Louisiana, Maine, Mississippi, Montana, North Dakota, Nebraska, New Hampshire, New Mexico, Nevada, Oklahoma, Rhode Island, South Carolina, South Dakota, Vermont, West Virginia, Wyoming and Puerto Rico. CoBRE grants are awarded in three 5-year phases. Data from CoBRE awards were retrieved using NIH Reporter by searching for the CoBRE Request for Applications (RFAs)/Program Announcements (PARs) (https://reporter.nih.gov/). Data for CoBRE awards from FY00 through FY22 were collected. There are different RFAs/PARs for the Phase I, Phase II and Phase III CoBRE grants and the data was initially collected in these three categories. The Phase I/Phase II awards are funded using the P20 grant mechanism and the Phase III awards are funded using the P30 grant mechanism. Initially, CoBRE awards were administered by the National Center for Research Resources (NCRR), with appropriate project numbers (e.g. P20RR020173). In late 2011, when the NCRR was eliminated, the National Institute of General Medical Sciences (NIGMS) took over administration of the IDeA program and CoBRE grants received GM project numbers (e.g. P20GM103464). For these reasons, each CoBRE grant might have two or three different grant numbers during the lifetime of the award. Different grant numbers for a single CoBRE were linked by comparison of the institution, title of the grant and/or the principal investigator of the grant. The data for each CoBRE contained all of the sub-projects affiliated with the CoBRE, with some exceptions. The administration, alteration and renovation, and research cores for each CoBRE were identified by the title of the sub-project. All other sub-projects, with the name of a research project rather than a core, were collected as research projects. The dataset is incomplete since research cores were not available as sub-projects prior to FY04 and CoBRE awards transferred from NCRR to NIGMS in FY12 did not have sub-projects available after the transfer. The transfer of CoBRE awards to NIGMS impacted the data from 13 Phase I, 32 Phase II and 15 Phase III CoBRE grants. The publications and patents associated with each CoBRE were collected using NIH Reporter on October 23, 2022.

Subsequent grants received by the project and pilot project leaders were identified by matching the Contact PI Person ID (PIID) for the CoBRE investigator with NIH R series grants. Some CoBRE investigators received R series awards prior to their start date on the CoBRE award. CoBRE investigators with prior R01 or R35 (NIH MIRA award) grants were categorized as established investigators, while investigators that did not have prior R01/R35 funding were categorized as new investigators. The analysis of investigator success focuses on the latter group of investigators.

R01 data from 21 different institutes was collected from NIH Reporter: the National Center for Complementary and Integrative Health (NCCIH), the National Cancer Institute (NCI), the National Eye Institute (NEI), the National Human Genome Research Institute (NHGRI), the National Heart, Lung and Blood Institute (NHLBI), the National Institute of Aging (NIA), the National Institute on Alcohol Abuse and Alcoholism (NIAAA), the National Institute of Allergy and Infectious Diseases (NIAID), the National Institute of Arthritis and Musculoskeletal and Skin Diseases (NIAMS), the National Institute of Biomedical Imaging and Bioengineering (NIBIB), the National Institute of Child Health and Human Development (NICHD), the National Institute on Drug Abuse (NIDA), the National Institute on Deafness and Other Communication Disorders (NIDCD), the National Institute of Dental and Craniofacial Research (NIDCR), the National Institute of Diabetes and Digestive and Kidney Diseases (NIDDK), the National Institute of Environmental Health Sciences (NIEHS), the National Institute of General Medical Sciences (NIGMS), the National Institute of Mental Health (NIMH), the National Institute on Minority Health and Health Disparities (NIMHD), the National Institute of Neurological Disorders and Stroke (NINDS) and the National Institute of Nursing Research (NINR). Data for other R series grants was collected from all institutes.

### Building comparison cohorts

One goal of the CoBRE grants is mentoring junior investigators to independence. A comparable NIH program is the K01 grants program. For comparative purposes a cohort of K01 awardees was analyzed. K01 awards with the first year of support between 2000 and 2017 were chosen for analysis. These are contemporaneous with the CoBRE awards. The cutoff of 2017 was chosen since the maximum duration of these awards is 5 years and thus were completed by FY22, which was the end date of the analysis. K01 awardees who subsequently received R01 funding were determined by matching the PIID between the K01 awardees and R01 awardees. A second goal of the CoBRE awards is the development of research infrastructure. The output of this investment was measured as research publications and patents. For comparison of the movement of successful CoBRE investigators to other institutions, a cohort of non-IDeA new investigators was identified. NIH trainees supported by F31, F32, K99 and K01 awards from FY01 through FY19 were collected using NIH Reporter. The PIIDs of this cohort were used to identify which of these trainees were awarded R01 grants. These PIs were further narrowed to exclude any PIs who held their R01 at an IDeA institution during year 1 of the award. This cohort was used for the comparison. The location of the award in year 1 was compared with the location of the award in year 4 to determine if newly funded investigators in the non-IDeA cohort took the opportunity to move to another institution.

### Other Data Sources

Research and Development expenditures at academic institutions was retrieved from NSF/NCSES and come from the Higher Education Research and Development survey (https://www.nsf.gov/statistics/srvyherd-legacy/). Data of interest included sources of funds for research and development expenditures (federal, state, institutional, business/industry and other) and federal agencies providing funds for research and development expenditures (DOD, DOE, HHS, NASA, NSF, USDA, others). For some years (2000–2009), several university systems reported their data at the system level, rather than the campus level. In these cases, the data for expenditures for each campus was estimated based upon the distribution of expenditures between individual campuses in other years (2010–2021).

### Activity codes and RFAs

NIH grants were categorized as research project, construction, capacity building, cooperative, outreach, training, infrastructure, equipment, center and resource grants and cooperative agreements in different analyses. NIH grant activity codes were used for categorization (https://grants.nih.gov/grants/funding/ac_search_results.htm).

### Statistical analyses

Outliers were identified using the IQR method using a threshold of 1.5 times the interquartile range. Graphical and statistical analyses were performed using GraphPad Prism. Simple linear regression was performed to measure the correlation between two variables. Fisher’s exact test was used to compare the distribution of two cohorts (e.g. CoBRE project leaders vs K01 awardees) between different categories (e.g. R01 funded vs not R01 funded). Multiple linear regression (least squares method) was used to examine the relationship between different sources of investment for research and development and NIH funding for research projects. The F statistic for the regression model was used to determine if the model predicted the dependent variable better than random. The goodness of fit of the model was estimated from R^2^. The D’Agostino Pearson test was performed to determine if the distribution of residuals was Gaussian. The absolute value of the t statistic (|t|) was calculated to determine if the effect of each variable on the model was greater than zero. Multicollinearity was determined by calculating R^2^ with the other variables. The introduction of more than one NIH funding variable, e.g. NIH research projects and CoBRE funding or NIH all other funding, increased multicollinearity in the models.

### WVU IRB Approval

The West Virginia University Institutional Review Board approved the study (WVU Protocol#: 2304764418).

## Results

### NIH Investment

The Center of Biomedical Research Excellence (CoBRE) grant program was initiated in 2000 as part of the Institutional Development Award (IDeA) program, which aims to geographically diversify NIH funding. Phase I of a CoBRE grant is for 5 years and supports research cores to develop and provide research infrastructure at IDeA institutions, and funds for junior faculty to develop research projects that are competitive for independent NIH funding. Each CoBRE has an administrative core and an important component of this core is a mentoring program to guide the development of junior faculty. CoBREs also support pilot projects and some have had a renovations core. A CoBRE can be renewed for a second 5-year period as Phase II, which is similar in structure to Phase I, and can be renewed for Phase III with an additional 5 years of support primarily to support research cores and some pilot projects. NIH has supported 209 Phase I CoBREs, 137 Phase II CoBREs and 77 Phase III CoBREs through FY22 (Table 1). These grants were held at 73 different academic/research institutions in 23 states and Puerto Rico (Supplemental Figure 1).

**Table 1.** CoBRE Grants Awarded by NIH.

|  | **Awarded** | **Completed** | **Active** |
| --- | --- | --- | --- |
| **Phase I** | 209 | 153 | 56 |
| **Phase II** | 137 | 90 | 47 |
| **Phase III** | 77 | 59 | 18 |

Awarded, completed and active CoBRE grants through the end of FY22.

The NIH investment in CoBRE grants (total costs) over time is illustrated in Figure 1A. There were two waves of investment in CoBRE Phase I grants, followed by waves of Phase II grants. The first wave of Phase II CoBRE grants was followed by a wave of Phase III grants. The average total costs awarded per Phase I grant (excluding supplements) was $2.08 million +/- $290,000 per year. Phase II CoBREs averaged $2.11 million +/- $201,000 and Phase III grants averaged $1.09 million +/- $80,000 per year. Changes in the average total cost per CoBRE grant per fiscal year is shown in Figure 1B. Simple linear regression analysis indicates that the average total cost per Phase I and Phase II CoBRE grants per fiscal year increased over time. Average total costs per Phase III CoBRE grants have not significantly risen over time.

**Figure 1.**
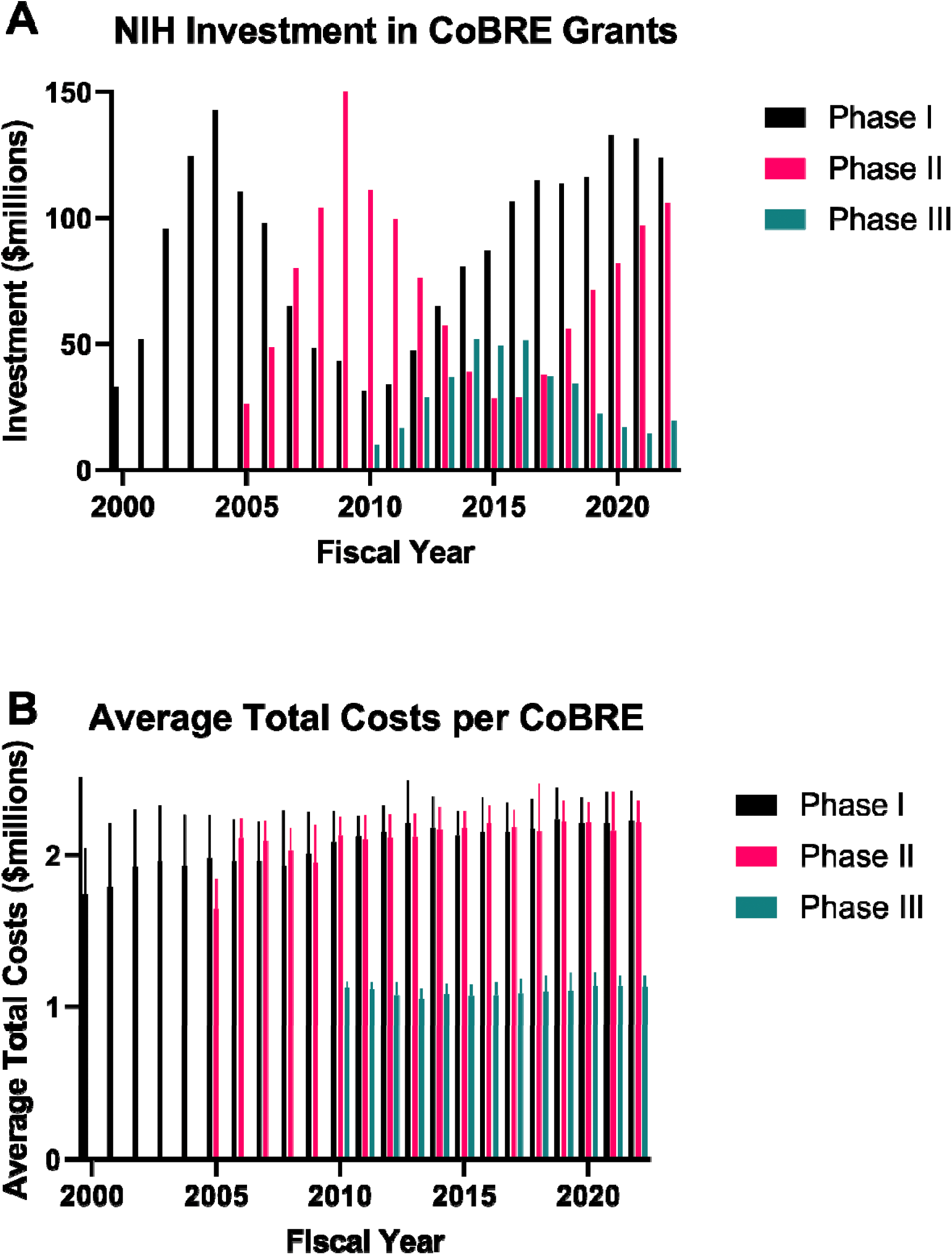
NIH Investment in CoBRE Grants. **A)** The total costs for all Phase I, Phase II and Phase III CoBRE grants in each fiscal year are illustrated. **B)** The total average costs of Phase I, Phase II and Phase III CoBRE grants in each fiscal year are shown.

One goal of the CoBRE grants is investment in research core infrastructure. The estimated investment in research cores in the CoBREs are illustrated in Figure 2. Prior to 2004, the costs for the research cores are not broken out as sub-projects and the data is unavailable. Sub-projects are unavailable in NIH Reporter for CoBRE awards that were transferred from NCRR to NIGMS in FY12. The investments in research cores in these CoBREs from FY12 through FY16 is estimated from the investment made in FY11. The total costs associated with the research cores for each Phase of CoBRE grant per fiscal year is shown. The summation of the total investment in research cores across all three phases of CoBRE grants per fiscal year is also shown. The total investment in research cores was relatively flat initially, but has seen a sustained increase over the last decade. A compilation of CoBRE-supported research cores extracted from NIH Reporter is shown in Supplemental Table 1.

**Figure 2.**
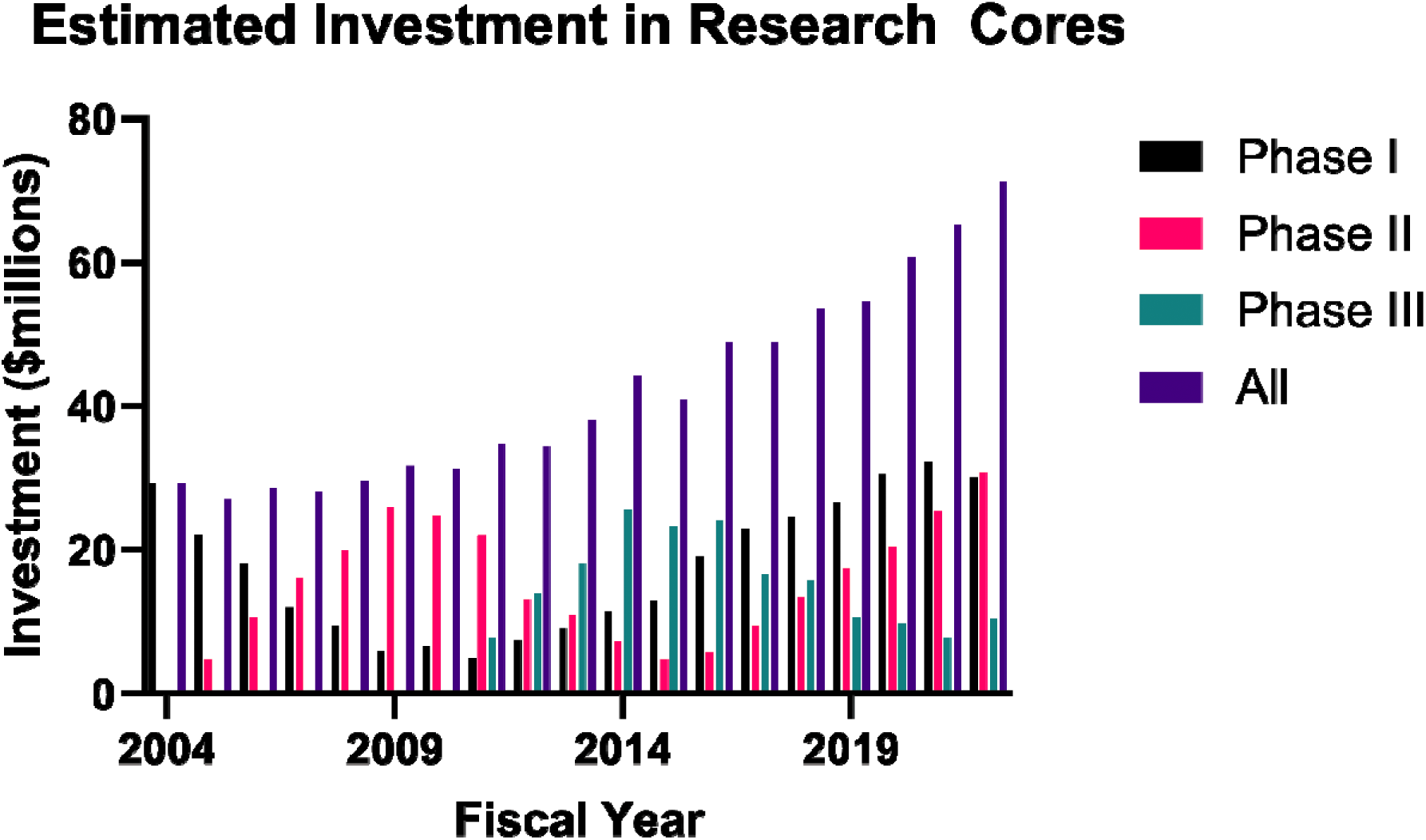
NIH Investment in CoBRE Research Cores. The total costs for all research cores in Phase I, Phase II and Phase III CoBRE grants in each fiscal year is shown. The summed total costs for all research cores (All = Phase I + Phase II + Phase III) per fiscal year is also shown.

A second goal of the CoBRE grants program is investment in the development of junior scientists by supporting research projects and pilot projects. Individual research projects affiliated with each CoBRE grant were identified based upon the sub projects. A total of 2070 different potential research project leaders were identified. Of these investigators, 158 were the principal investigator on an R01 or R35 in a fiscal year prior to their first year of support on the CoBRE grant. The remaining 1912 investigators were new and/or early-stage investigators without prior major funding from the NIH, although some of these new investigators had prior NIH support in the form of R21 or R15 grants. Support for these individuals ranged from a few thousand dollars to the bulk of the entire CoBRE budget. Since the identification of project leaders is imprecise, outliers were excluded to restrict the analysis to individuals with meaningful and realistic support for an individual CoBRE research project. There were no outliers identified at the lower end of projects supported in Phase II, so an arbitrary cutoff of a minimum of $10,000 was used. Exclusion of outliers provided a cohort of 1728 investigators for the analysis. The total investment in these Phase I research projects (total costs) was $694,123,823 and in these Phase II projects (total costs) totaled $462,531,134. The costs per project per fiscal year and the total investment in individual investigators over the course of the CoBRE are shown in Figure 3. The average support for a research project per fiscal year was $222,476 +/- $67,824 in Phase I and $190,342 +/- $82,679 for Phase II. The average total support over the course of the CoBRE per investigator in Phase I was $597,192 +/- $410,812 and in Phase II was $441,864 +/- $345,783.

**Figure 3.**
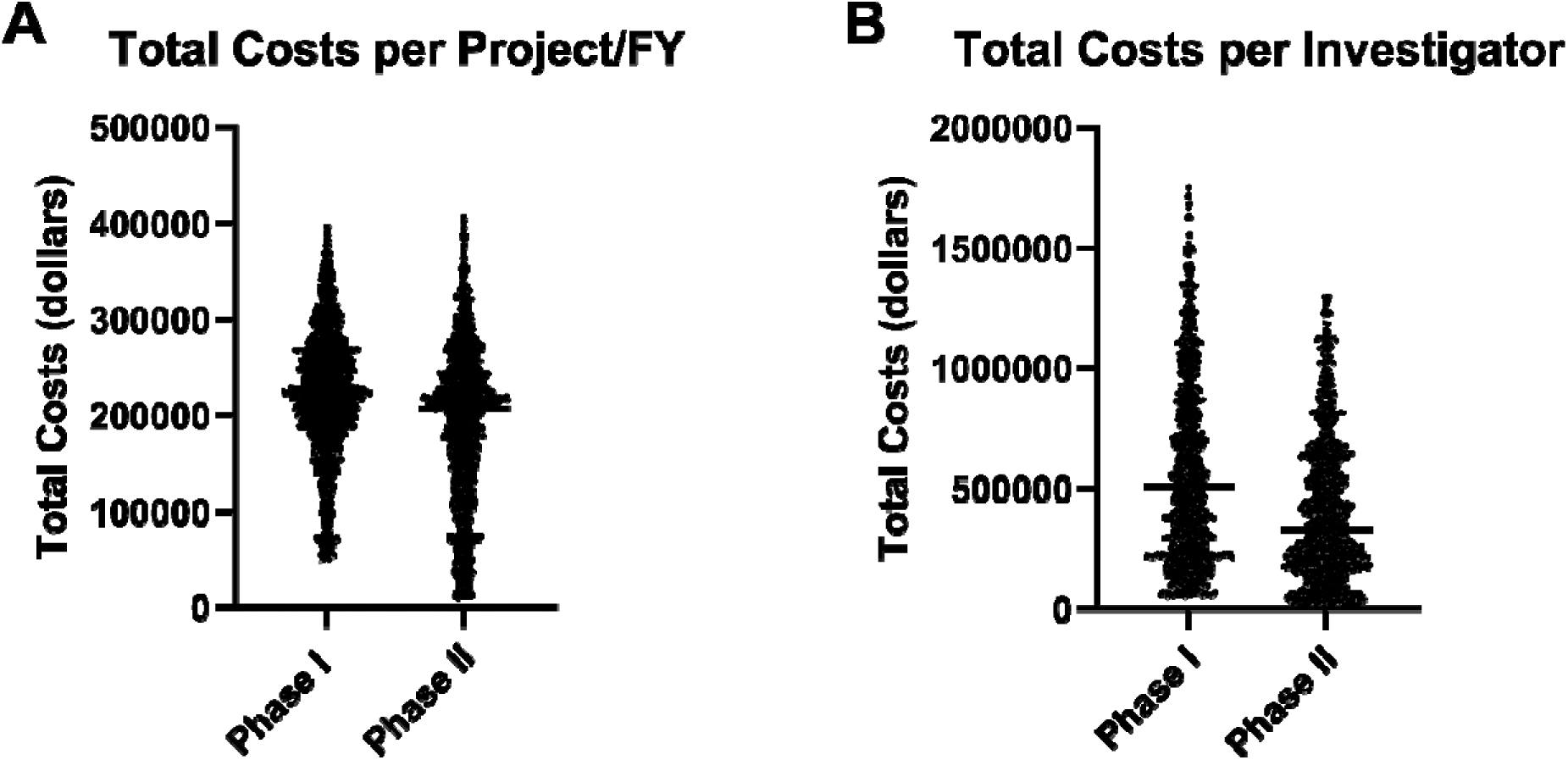
Investment in CoBRE Research Projects. **A)** The total costs per fiscal year, FY00 through FY22, for each project supported by Phase I CoBREs and Phase II CoBREs, excluding outliers. Each data point reflects a single project in one fiscal year. Projects supported for multiple years have multiple data points. **B)** The total costs provided to each investigator, FY00 through FY22, for Phase I CoBREs and Phase II CoBREs, excluding outliers. Each data point represents a single investigator. Note that one investigator can be supported in Phase I and Phase II of a CoBRE. For these individuals the costs incurred in Phase I are reflected in the left plot in each panel and those incurred in Phase II are reflected in the right plot in each panel.

### CoBRE Outcomes – Grants

An essential outcome for the success of CoBRE grants is the transition of junior investigators to independent research funding. This analysis focuses on NIH funding for two reasons. First, the mission of the IDeA program is to geographically broaden the distribution of NIH research dollars. Second, NIH data is robust and new investigators on CoBRE grants can be linked to subsequent R series grants, which cannot be done for other federal agencies or foundations. Thus, this approach *underestimates* grant success by limiting the measurement to NIH R series grants and misses project leaders who successfully secure funding from other federal agencies and foundations.

A total of 740 new investigators were subsequently awarded an R series grant, i.e. an R01, R35, R21, R15, R41, R42, R43 or R44 grant (Table 2). Thus, 39.2% of the CoBRE project leads were awarded an R series grant. These investigators were awarded 1391 R series grants that had total costs of $1,638,867,232 over the course of their careers (Table 3). Major awards, an R01 or R35, were secured by 551 new investigators. These investigators were awarded 888 major awards that had total costs of $1,435,118,545. Thus, 31.9% of new investigators received a major award subsequent to their support on a CoBRE grant. Of these investigators, 83.8% received a major award within 5 years of their initial support on a CoBRE grant. These numbers underestimate the success of CoBRE investigators, since the majority of CoBRE grants are still in Phase I or Phase II. If the analysis is restricted to the investigators from the 90 Phase I/Phase II CoBREs that are completed, 40% of the investigators secured an R01 or R35.

**Table 2.** Number of New Investigators Receiving R Series Grants.

|  | **R01s** | **Only**<br>**R35s** | **Only**<br>**R21s** | **Only**<br>**R15s** | **Only**<br>**R41/R42** | **Only**<br>**R43/R44** | **Total** |
| --- | --- | --- | --- | --- | --- | --- | --- |
| **# Invest.** | 531 | 20 | 77 | 44 | 3 | 2 | 677 |

The number of unique investigators receiving R series grants. 531 investigators were awarded R01s, 31 were awarded R35s, but 11 of the R35 awardees had already received an R01. Therefore 20 investigators received only an R35. In a similar fashion, the table shows the number of unique investigators securing only an R21, R15 and R40 series grants.

**Table 3.** Number of R Series Grants Awarded to New Investigators.

| **Grant** | **Number Investigators** | **Number of Grants** | **Total Costs** |
| --- | --- | --- | --- |
| **R01s** | 531 | 857 | \$1,396,755,719 |
| **R35s** | 31 | 31 | \$38,362,826 |
| **R21s** | 258 | 403 | \$156,772,117 |
| **R15s** | 56 | 69 | \$28,770,517 |
| **R41/R42** | 19 | 28 | \$15,410,045 |
| **R43/R44** | 3 | 3 | \$2,796,008 |
| **TOTAL R Series Grants** | | 1391 | \$1,638,867,232 |

Note that every grant is counted once. A single investigator can appear multiple times in the table, e.g., if they are awarded an R01, an R35 and an R21 the investigator will appear in row 1, row 2 and row 3.

**Table 4.**
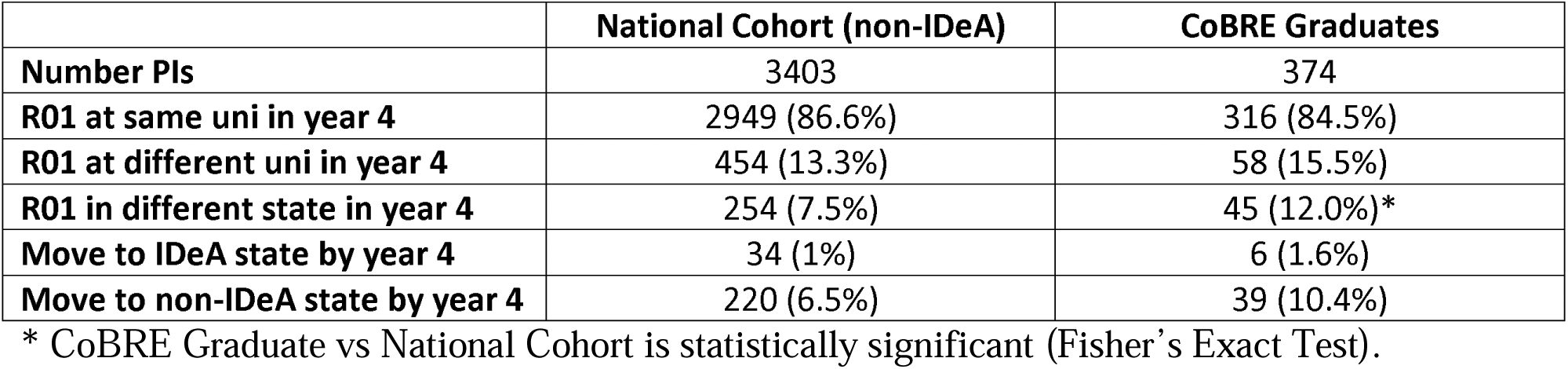
Movement of Investigators After Securing R01 Funding.

|  | **National Cohort (non-IDeA)** | **CoBRE Graduates** |
| --- | --- | --- |
| **Number PIs** | 3403 | 374 |
| **R01 at same uni in year 4** | 2949 (86.6%) | 316 (84.5%) |
| **R01 at different uni in year 4** | 454 (13.3%) | 58 (15.5%) |
| **R01 in different state in year 4** | 254 (7.5%) | 45 (12.0%)* |
| **Move to IDeA state by year 4** | 34 (1%) | 6 (1.6%) |
| **Move to non-IDeA state by year 4** | 220 (6.5%) | 39 (10.4%) |

\* CoBRE Graduate vs National Cohort is statistically significant (Fisher's Exact Test).

The K01 award is a Mentored Research Scientist Development Award that supports a junior scientist’s development toward independence. The goal is similar to one of the goals of the CoBRE programs. For comparison to the CoBRE awards, new K01 awardees from FY00 through FY17 were identified using NIH Reporter. The K01 award is up to 5 years and selection of this time frame meant that these awards ended in FY22 at the latest, i.e. all the awardees analyzed had completed their K award. This cohort consisted of 3283 K01 awardees and 42.5% of these awardees successfully secured an R01. In IDeA states, 40% of K01 awardees (215 individuals) successfully transitioned to an R01. The difference in number of K01 awardees getting an R01 in IDeA states was not significantly different than those in all other states. Comparison with the 90 CoBREs that completed Phase II, 40% of new CoBRE investigators received an R01. The successful transition of K01 awardees to R01 awardees is statistically the same as the rate for CoBRE investigators (Fisher’s exact test).

Progression from CoBRE support to independent support was measured by the time required to secure a major NIH award. The distribution of time until R01/R35 was examined in two ways. From the perspective of the CoBRE support, the time from the start date of the CoBRE grant (Phase I) to the start date of the first R01/R35 awarded to the investigator was measured (Figure 4). The elongated distribution over time is expected, since an investigator beginning support on the Phase II CoBRE in year 3 (year 8 overall) might be awarded an R01 in year 5 of the Phase II award (year 10 of the CoBRE overall). This analysis reports the distribution of new major awards to investigators over the course of the CoBRE grant and beyond. The time from the initial start date of each investigator on the CoBRE to the start date of the subsequent R01/R35 was also measured (Figure 4). This analysis more meaningfully measures the progression of each successful investigator to an independent major award. The median time to securing a major award is 3 years from beginning support on the CoBRE (mean = 3.2 years +/- 2.89 years) and the 75^th^ percentile is 4 years.

**Figure 4.**
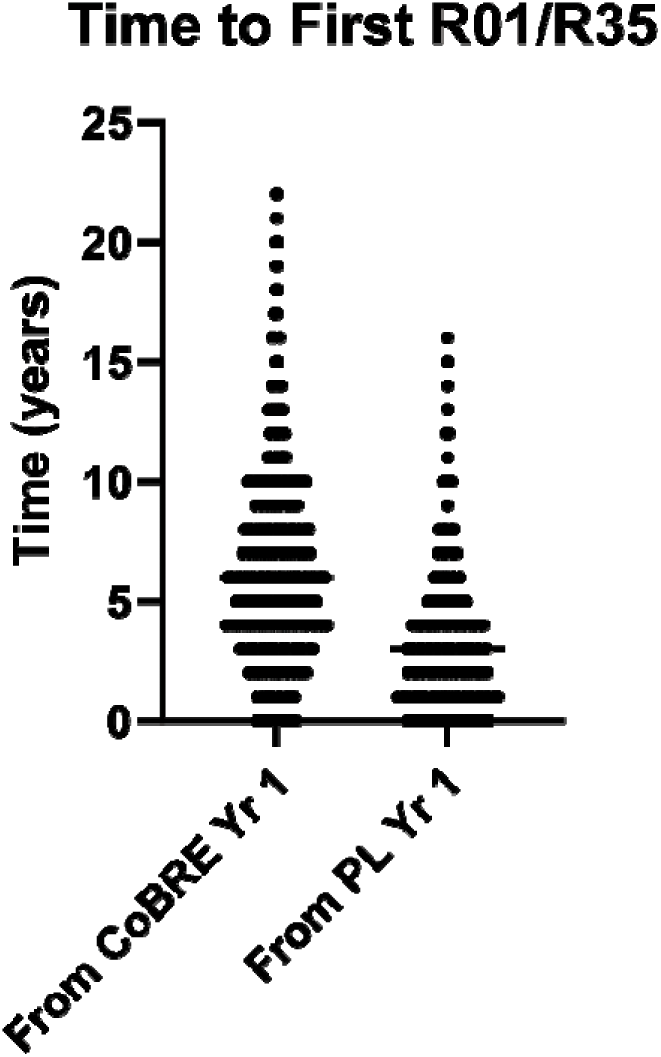
Time to First R01/R35. The time in years until each new investigator supported by a CoBRE grant receives their first R01 or R35 grant is illustrated. The data on the left measures the time from the start date of the CoBRE grant (Phase I) until new investigators receive a major award and reflects awards made relative to the lifetime of the CoBRE. The data on the right measures the time between a new investigator’s support on a CoBRE and the start date of a subsequent R01 or R35. Each data point represents a single investigator.

The number of major awards and all R series grants awarded to CoBRE investigators is shown in Figure 5. The average number of R01/R35s per CoBRE is 4.4 +/- 5.0 with the most successful CoBRE mentoring investigators who were awarded 23 major NIH grants. The average number of R series grants was 6.7 +/- 7.4 per CoBRE, with a maximum of 48 total grants for one CoBRE. Since more than half of the total CoBREs are still active, this is an underestimate of success. Restricting the analysis to CoBREs that have completed Phase II reveals an average of 7.4 +/- 5.8 R01/R35 grants and 11.5 +/- 8.3 total R series awards.

**Figure 5.**
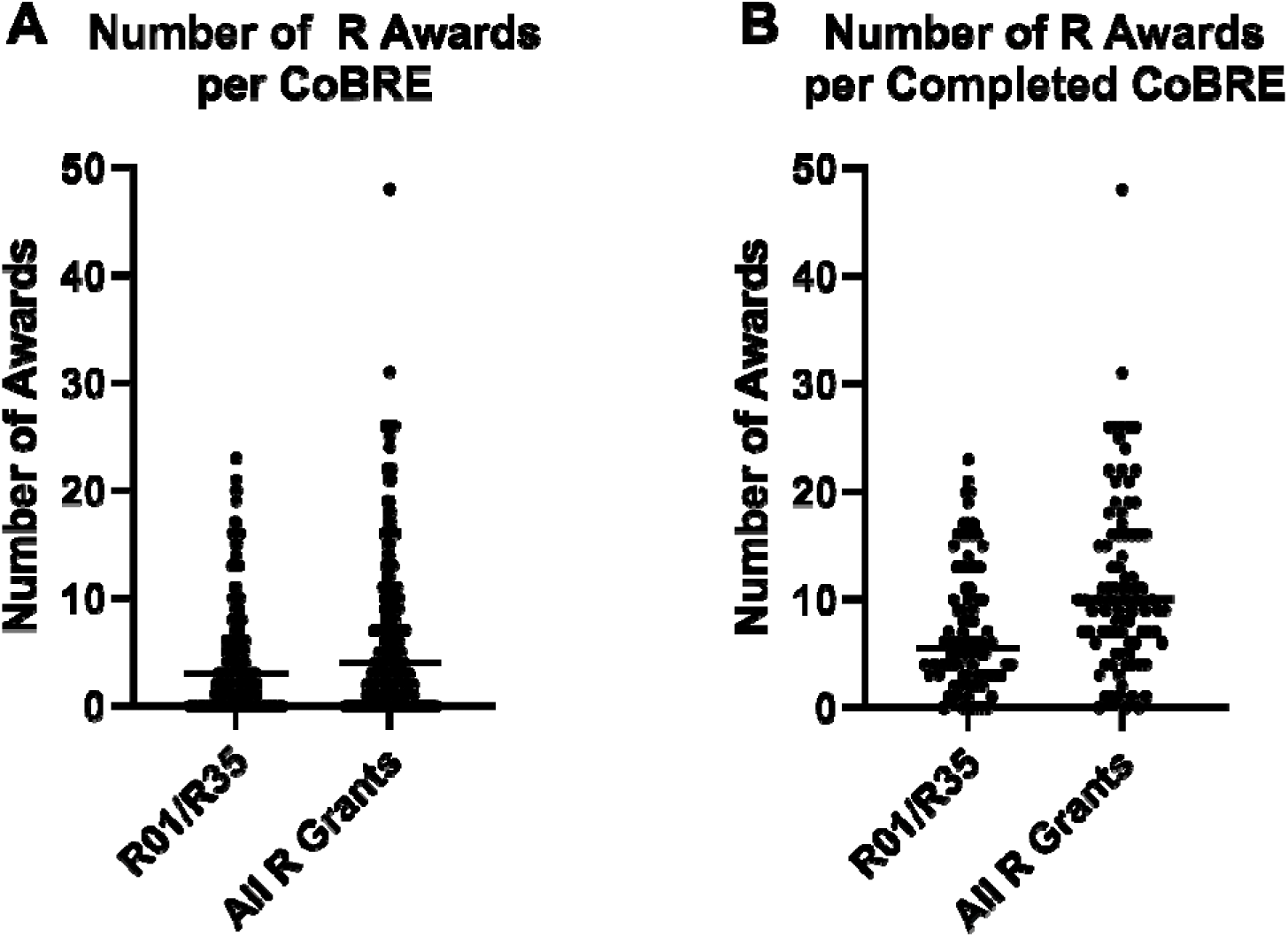
The Number of R series Grants Awarded to New Investigators Supported on CoBRE Grants. The total number of all R01/R35s or all R series grants that were awarded to new investigators supported by CoBRE grants are illustrated. The data for all CoBREs (both active and completed) is shown in the left panel (**A**) and the data for all CoBREs that completed Phase II is shown in the right panel (**B**). Each data point represents a single CoBRE award.

### CoBRE Outcomes – Publications and Patents

Based upon data extracted from NIH reporter, 30,800 unique papers affiliated with CoBRE grants were published as of this analysis. There were 26,213 papers linked to the Phase I/Phase II CoBRE grants and 4,587 papers from the Phase III CoBREs. The number of papers published per CoBRE ranged from 0 to 665 with a median of 130 papers (see Figure 6). The time from the CoBRE start date to the date of each publication (in years) was calculated and plotted. Seventy-eight percent of the papers associated with CoBRE grants are published within 11 years of the start date of the CoBRE (Phase I). CoBRE investigators have filed 123 patents.

**Figure 6.**
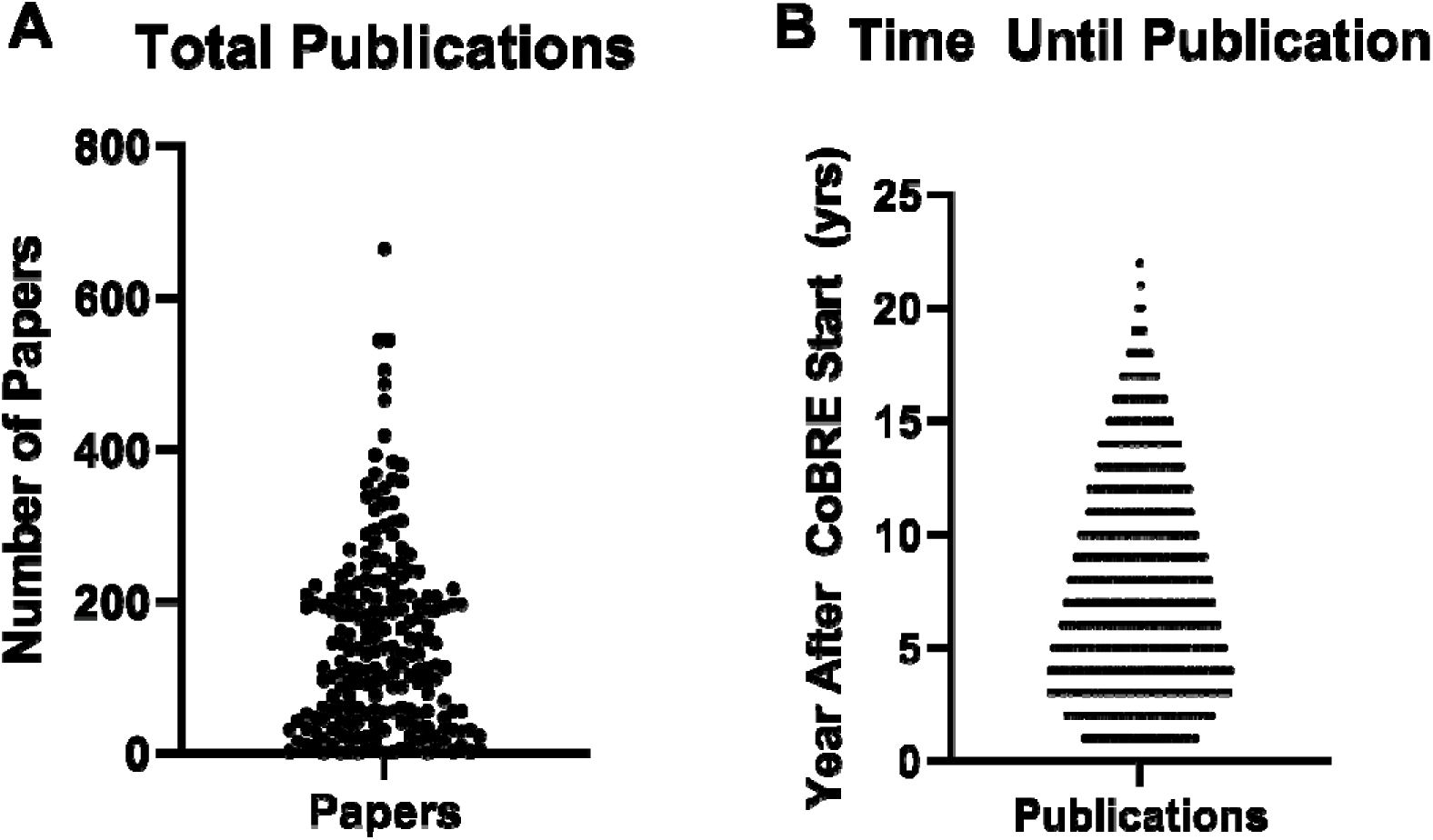
Number of Publications and Elapsed Time Until Publication. The total number of publications affiliated with each CoBRE grant is illustrated on the left (**A**). Each data point represents a single CoBRE grant. The time between the start of Phase I of a CoBRE grant and the year of publication is illustrated in the panel on the right (**B**). Each datapoint represents a single publication.

### Migration of CoBRE Investigators

One concern with the CoBRE mechanism of support is that junior investigators will graduate from the CoBRE by securing major NIH awards and then move to another institution. To address this issue, investigators awarded an R01 that has gone into year 4 were analyzed to compare the institution where the R01 was held in year 1 and year 4 (Table 5). Of the 403 PIs analyzed, 16.1% had changed institutions between year 1 and year 4 of their R01 and 10.7% moved to a non-IDeA state. A cohort was established to compare with the CoBRE graduates. NIH trainees on F31, F32, K99 and K01 awards from FY01 through FY19 were captured. Trainees who were subsequently awarded an R01 that was held at an institution in a non-IDeA state in year 1 were compiled. This cohort of 3,403 PIs received their first R01s contemporaneously with the CoBRE graduates. The two cohorts were compared using Fisher’s exact test. The percentage of CoBRE graduates moving to another institution between year 1 and 4 was not significantly different (p = 0.26) from the percentage of the non-IDeA cohort. While the percentage of CoBRE graduates moving to a different state was significantly more than the non-IDeA cohort (p = 0.0017), the percentage of PIs moving to IDeA vs non-IDeA states was not significantly different between the two cohorts (p > 0.9999). These findings demonstrate that investigators supported by the CoBRE mechanism are no more likely than their counterparts in non-IDeA states to change institutions upon securing a major NIH award.

**Table 5.**
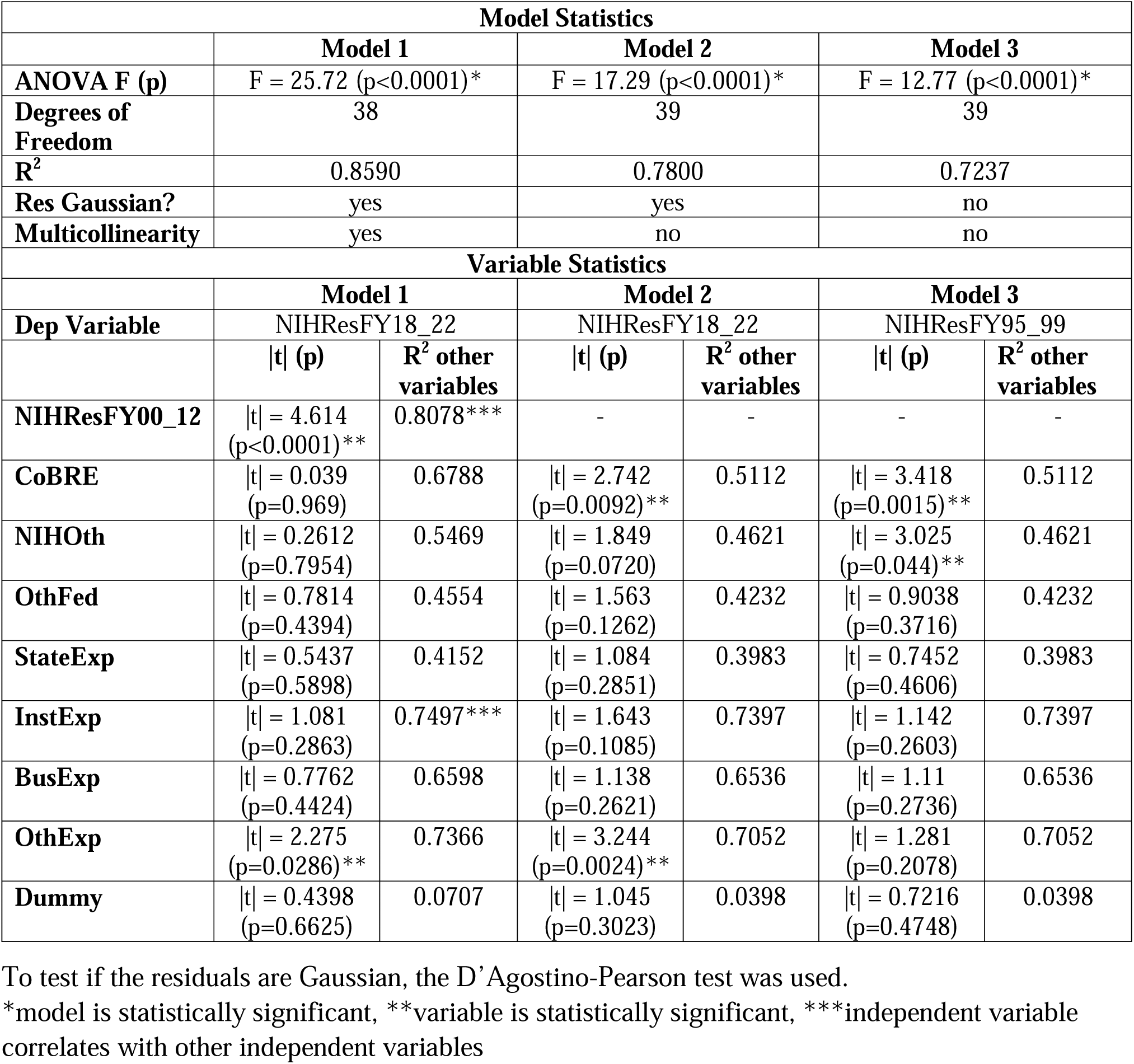
Statistical Analysis of Mathematical Models.

### CoBRE Impact

A prediction of successfully building research capacity and increasing competitiveness for NIH funding in IDeA states is that these states would increase their share in NIH research funding. Historically, the number of R01s awarded in IDeA states paralleled the overall changes in funding by the NIH (see supplemental Figure 1). In 2000, concomitant with implementation of the CoBRE program, the proportion of NIH funding to IDeA states increased (see Figure 7A). The increased share of NIH funding was sustained until 2016, which coincided with end of the flatline budget at NIH and the beginning of annual increases to the NIH budget. This trend in funding is generally reflected in the expenditures of federal R&D funding at IDeA institutions from the HERD survey.

**Figure 7.**
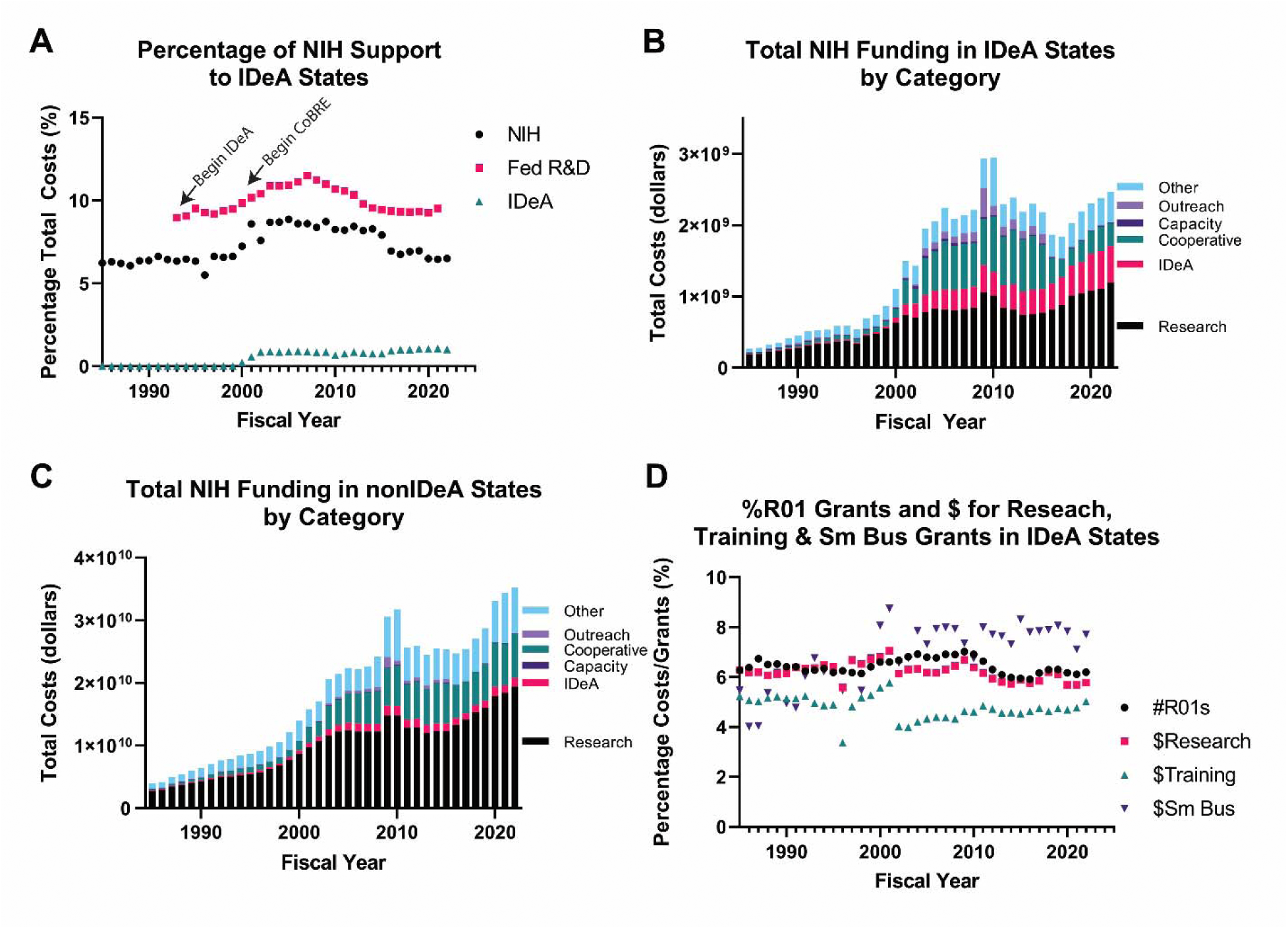
Historical Changes in NIH Funding to Recipients in IDeA States. **A)** The percentage of total NIH funding invested in IDeA states for each fiscal year is shown (since 1985). The percentage of total expenditures of federal R&D funds at academic institutions in IDeA states, as per the HERD survey, is also shown (FY91 through FY21 – based upon data availability). Also shown is the percentage of total NIH funds that are awarded to IDeA states through the INBRE, CoBRE and IDeA-CTR programs. **B)** The total costs of NIH awards received in the IDeA states in each of six categories. Note the increase after 2000 in the IDeA category (P20, P30 and U54 grant mechanisms, which are used for the INBRE, CoBRE and IDeA-CTR programs), the cooperative grants and outreach/intervention categories. **C)** The total costs of NIH awards received in the non-IDeA states in each of six categories. While these states are ineligible for IDeA program funding, NIH uses P20, P30 and U54 mechanisms to fund other types of grants. Note the increase in funds in the cooperative grant and outreach/intervention categories, although the increases are less than the increases observed in the IDeA states. **D)** The percentage of total R01s funded that were awarded in IDeA states for each fiscal year is shown. The percentage of total NIH funds in research grants, total NIH funds in training grants and total NIH funds in small business grants awarded in IDeA states is also shown.

The increase in NIH funding in IDeA states came from several funding mechanisms (Figure 7B). First, there was an increase in funding via the grant mechanisms that support INBRE (P20), CoBRE (P20 and P30) and IDeA-CTR (U54) programs. Second, there was an increase in cooperative grants awarded to IDeA states (U series of grants). Third, there was an increase in grants in the areas of outreach and intervention in the IDeA states. There was also an increase in cooperative grants and outreach/intervention grants awarded to non-IDeA states, but they had a larger impact on funding to IDeA states (Figure 7C). The proportion of R01s funded in IDeA states is relatively unchanged since 1985 (Figure 7D). Similarly, the proportion of NIH funds awarded for all research grants in IDeA states has been constant. The proportion of NIH funds awarded for training in IDeA states dropped from approximately 6.2% in FY01 to 4.3% in FY02. While on an upward trajectory, the proportion of NIH training funds awarded in IDeA states in FY22 was only 4.9%.

The impact of CoBRE grants is expected at the institution level rather than the state level. The total costs of CoBRE grants at 48 academic institutions was compared with the total costs of NIH research grants from FY18 through FY22 at those institutions by simple linear regression and r squared was 0.5264. The analysis was extended to examine the relationship between various research and development investments using a mathematical model to correlate with the total costs received from NIH for research projects from FY18 to FY22. The data was fitted using the equation:

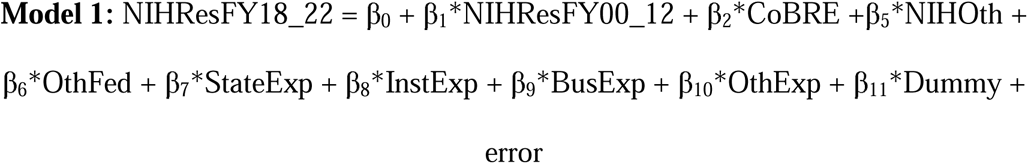

The variables in the model are:

NIHResFY18_22 - the sum of total costs of NIH awards for research projects at the institution from FY18 to FY22

NIHResFY00_12 - the sum of total costs of NIH awards for research projects at the institution from FY00 through FY12

CoBRE - the sum of total costs of all CoBRE awards to the institution

NIHOth – the sum of total costs of all NIH awards for construction, infrastructure, equipment, capacity, center and resource and cooperative agreements at the institution

OthFed - the sum of all R&D obligations from federal agencies other than the HHS (parent agency of NIH) from FY00 through FY21 at the institution

StateExp - the sum of all R&D expenditures of state funds at the institution from FY00 through FY21

InstExp - the sum of all R&D expenditures of institutional funds at the institution from FY00 through FY21

BusExp - the sum of all R&D expenditures of business/industry funds at the institution from FY00 through FY21

OthExp - the sum of all R&D expenditures of all other funds, including from non-profit organizations, at the institution from FY00 through FY21

Dummy – a dummy numeric value based upon the first letter of the institution (e.g. A = 1, B = 2, etc.)

The relationship of these variables with the total costs of NIH awards for research projects in FY18 to FY22 was determined by multiple regression analysis. Two variables correlated with funding for research grants from FY18 to FY22, previous funding for research grants from NIH (from FY00 to FY12) and R&D expenditures from other funds, including non-profit organizations (Table 5). The model was revised to eliminate the variable NIHResFY00_12 to determine if other variables correlated with NIHResFY18_22 in the absence of the dominant variable. Multiple linear regression demonstrated a significant relationship between CoBRE funding and R&D expenditures from other sources and NIHResFY18_22 (Table 5 – Model 2). The model was revised to replace NIHResFY18_22 with NIHResFY95_99, i.e. to compare NIH research grant funding prior to implementation of the CoBRE funding mechanism with CoBRE funding. The two variables correlated with NIHResFY95_99 were CoBRE funding and other types of funding from the NIH (Table 5 – Model 3). Since CoBRE funding correlated with NIH research grant funding prior to CoBRE awards, the relationship likely indicates that institutions that can attract NIH research grants are also more successful at securing CoBRE funding.

To examine the relationship of changes in funding of research projects temporally relative to CoBRE funding, 21 academic and research institutions with multiple CoBRE grants awarded between 2000 and 2006 were analyzed. Research project funding over a 20-year time period, from 5 years before the start data of the first CoBRE (yr = -5) to the end of Phase III of the first CoBRE (yr = 15). NIH research project funding at the institution as a percentage of total NIH research project funding for each FY was calculated. The percentage of NIH research project funding at the institution in year -5 was normalized to 1. The data is shown in supplemental figure 3. The results are mixed. Approximately a dozen institutions showed an increased percentage of NIH research funding after receiving the CoBRE award, whereas others did not.

## Discussion

Cornerstones of the CoBRE programs are support for the research programs of junior investigators and mentoring programs to foster the development of these faculty into independently funded investigators. Approximately 31.4% of project leaders for all CoBREs successfully secured an R01/R35 grant. Including only the 90 CoBREs that completed Phase II, 40% of project leaders received an R01/R35 grant. A previous analysis of CoBRE grants examined the 19 CoBRE grants that were initially awarded (Phase I) in 2000 (14). A total of 107 junior investigators joined these CoBREs between 2001 and 2003 inclusive, and as of 2007, 40% were awarded an R01 grant (14). These data were collected on average 5.5 years after the investigators joined the CoBRE. This finding is consistent to the larger analysis here of 90 CoBREs completing Phase II. The impact of CoBRE activities upon junior investigator success was evaluated at the University of Nevada. Twenty junior investigators who had CoBRE support were compared with 20 junior investigators who were part of unsuccessful CoBRE applications, and therefore did not benefit from CoBRE activities. Success was defined as becoming PI on an extramural grant with a duration of at least 2 years and publishing an average of 1 paper per year. More of the CoBRE-supported investigators (47%) were successful than the investigators without CoBRE support (15%) (15). The outcomes of a CoBRE from Rhode Island were recently published where 6/10 project leaders and 5/18 pilot project leaders were awarded R01 grants (16). Thus, 39% of the pilot/project leaders secured R01 funding. The success rates of developing independently funding investigators across all of these studies are very consistent. These success rates are comparable to other NIH-supported mentored career awards (reported here and (14)). Collectively, these studies demonstrate the positive impact of CoBRE mentoring upon the development of junior investigators.

The impact of mentoring upon the success of early career scientists is established. The K-series of NIH grants are mentored career research awards. Two studies evaluated the impact of mentoring among a cohort of K-series applicants. The success of funded applicants in the cohort was compared with unfunded applicants. Both studies show that the funded (and mentored) applicants are more successful at securing additional funding than the unfunded (without structured mentoring) applicants (17, 18). Mentoring focused on grant writing at one institution has elevated the success rate of NIH R-series grant applications approximately 2.5-fold (19). A mentoring program for clinical faculty reports that a high percentage (92%) secure funding, although there is no comparative cohort in the analysis (20). The National Research Mentoring Network (NRMN) has multiple mentoring programs with different structures and each draws upon a national pool of junior investigators (21). The outcomes demonstrate that graduates of these mentoring programs are more successful at securing NIH funding than the national average (22, 23). These positive effects of mentoring strongly support the rationale for the mentoring component of the CoBRE grants.

Investment in CoBRE research core facilities has significantly impacted IDeA institutions. These facilities serve the needs of the project leaders, pilot project leaders and other researchers at the institution. Of the 30,800 publications linked to CoBRE grants in NIH Reporter, 80% are from projects that are not linked to the project leaders of the CoBREs. This reflects the utilization of CoBRE core facilities by researchers who are not directly supported by the CoBRE grant and underscores the impact of infrastructure building by these programs.

The CoBRE program is one component of the IDeA Program at NIH, which was created to build research capacity in states that were historically underfunded, with the goal of raising competitiveness for NIH funding and increasing the geographic distribution of research funding. In the 30 years of the IDeA Program, and the 23 years of the CoBRE program, the percentage of NIH funding, funding for NIH research projects, R01s and investment in training to IDeA states has not sustainably changed (Figure 7). While a mathematical model does demonstrate a correlation between CoBRE funding and NIH research funding, the relationship is not likely to be causal since there is a similar correlation between CoBRE funding and NIH research funding *prior* to the existence of the CoBRE program.

The EPSCoR programs at other federal agencies were also created to increase the geographic distribution of federal research funding, albeit the mechanics of the programs differ from the CoBRE mechanism. Evaluations of the success of the EPSCoR programs are mixed, but it is notable that “no state has ever ‘graduated’ from EPSCoR” (24–26). Analyses concur that the aggregate share of federal research dollars to EPSCoR states has not changed significantly. One analysis suggests that the EPSCoR programs contribute to the growth of federal research and development obligations at a rate of 0.0033% per year (26). A second analysis comparing the growth in federal research and development obligations between 2002 and 2005 suggests an increase in share of federal funding in a majority of EPSCoR states (57%) (24). Only 46% of non-EPSCoR states showed an increase in share of federal funding (24). Another analysis did not focus explicitly on state-by-state analysis, but rather focused upon cohorts based upon the first year of EPSCoR support, e.g. 1980 cohort, 1985 cohort etc. The analysis examined the change in percentage of NSF R&D funding per state in the cohort between the initial EPSCoR support and 2008. The results demonstrate an increase in the share of NSF funding for the early cohorts, but not for the later cohorts (27). In general, the effect of the EPSCoR programs upon federal funding is small. Evaluations and recommendations for the EPSCoR programs suggest that the goal of increasing the share of federal funding may be unattainable since non-EPSCoR institutions continue to invest in their research capacity, even as federal agencies specifically invest in the research capacity in EPSCoR states (25). However, the EPSCoR investment has resulted in significant positive effects at EPSCoR institutions including changes in research culture, expanding the research base, increasing research programs and elevation of research status (Carnegie Foundation rankings) (25, 27). A recent comparison of research output from institutions in EPSCoR and non-EPSCoR states also concludes that federal investments in EPSCoR states are effectively utilized (28). The gap in publications per faculty member at ESPCoR vs non-EPSCoR institutions has dramatically narrowed since 2011. Further, the number of publications per $1 million in federal research funding is substantially higher at EPSCoR institutions compared with non-EPSCoR institutions (28). Thus, the EPSCoR programs and the NIH CoBRE program have both significantly impacted individuals and institutions, but have not resulted in increased geographic distribution of research funding.

### Limitations

There are a number of limitations to this study. The dataset is incomplete and there are incidents of CoBRE awards lacking sub-project information, which is necessary for this analysis. Nevertheless, it is estimated that 92% of this information has been captured. Identification of project leaders on the CoBRE grants from NIH Reporter is imprecise. Roles are clearly defined on some CoBREs, but not the majority, and roles cannot be inferred based upon level of support since the range of financial support to individual projects is continuously distributed. As the range of support for projects ranged from a few thousand dollars, which is unlikely to impact the project, to the majority of the entire CoBRE budget, which are unlikely to be authentic project leaders, outliers were removed prior to the analysis. Measuring the success of project leaders based upon securing NIH grants underreports the success rate since grants from other federal agencies and foundations are excluded. The advantage of using NIH records is the ability to unequivocally link R series grants with project leaders using the Contact PI Person ID (PIID). Publications and patents associated with each CoBRE grant was extracted from NIH Reporter. There are errors in this information and the data was curated to remove records clearly not associated with the CoBRE, e.g. publications prior to the initiation of the CoBRE award. There is missing data in the HERD survey for institutions with relatively low levels of R&D expenditures (prior to 2005) and some data from the HERD survey required extrapolation from systems level data to campus level data.

### Concluding Remarks

Since the implementation of the CoBRE program there has not been an increase in geographic distribution of funding, i.e. increased proportional funding in their target states. The investment in the CoBRE program has been relatively small, e.g. approximately 1% of the NIH extramural budget in FY22. One potential solution is to increase the investment. There is evidence that the CoBRE grants are building research capacity by junior faculty securing independent extramural funding and the large number of papers and patents linked to the CoBRE grants (described above and (14)). There is evidence that other IDeA programs are having an impact, e.g. the IDeA-CTR grants are leading to clinical trials networks and improved service to their populations (29, 30). Based upon these successes, a second potential solution is to identify additional mechanisms to support research infrastructure complementary to the CoBRE grants. These grants might provide support for established investigators, rather than early-stage investigators, and include program-project and additional center like grants and could be supported in collaboration with other NIH Institutes. Another potential solution is to provide additional funding for training, e.g. the Leading Equity and Diversity in the Medical Scientist Training Program (LEAD MSTP) (31). There is evidence that the INBRE program is successfully impacting STEM students at the undergraduate level (32–36). Additional investment in graduate, postdoc and junior faculty training in the IDeA states is warranted given that the proportion of funding is below recent historic standards (Figure 7D). Further, federal support for graduate training lags behind the number of PhD students trained in these states (37). Since a larger proportion of PhD students trained in IDeA states hold their first R01s at institutions in IDeA states than students who train at other institutions, providing additional support for training at the graduate level might be an effective mechanism to build research capacity in IDeA states (37). A fourth solution is to examine institutions that have successfully increased their share of NIH funding, e.g. some of the institutions in supplemental Figure 3, and determine the factors leading to increased funding. These findings might establish best practices and provide a blue-print to replicate at other institutions that are historical underfunded by NIH.

## Author Contributions

The author was the sole contributor to the conceptualization, design, data collection, analysis and drafting the manuscript.

## Acknowledgements

Thanks to Drs. Pete Mathers, Karen Martin and Eric Kelley for insightful comments on the manuscript. MDS is the Director of the Cell & Molecular Biology and Biomedical Engineering Training Program (T32 GM133369) and the Director of Mentoring on the WVU Visual Sciences CoBRE (P20 GM144230).

## Data Availability Statement

Data are publicly available from NIH Reporter (https://reporter.nih.gov/) and from the NSF/NCSES Higher Education Research and Development survey (https://www.nsf.gov/statistics/srvyherd-legacy/).

---

**Supplemental Figure 1. Locations of Institutions Awarded CoBRE Grants.** The geographic locations of institutions that have been awarded CoBRE grants are shown. The figure is color-coded to illustrate the number of CoBRE grants that have been held by each institution.

**Supplemental Figure 2. Historical NIH Funding Since 1985.** The total costs of all awards from the NIH **(A)** and total number of R01s awarded by the NIH **(C)** plotted for each fiscal year. The total awards and awards to recipients in IDeA states are shown. The data for the IDeA states is plotted on a different scale in panels **(B)** and **(D)**.

**Supplemental Figure 3. Temporal Changes in Proportion of Research Project Funding Relative to CoBRE Start Dates at Selected Institutions.** Each academic/research institution and the years of the start dates of Phase I CoBREs at that institution (between 2000 and 2006) are indicated for each curve. The proportion of NIH research project funding (award activity codes including DP1, DP2, DP3, DP4, DP5, P01, R00, R01, R03, R04, R06, R15, R16, R19, R21, R22, R23, R24, R29, R33, R35, R27, R55, R56, R61, RM1, RF1, RC1, RL1 and RL2) at each institution was determined for a 20 year period, beginning 5 years before the start of the earliest CoBRE award at the institution (Year = - 5) and ending at Year = 15 (approximate end date of Phase III of the first CoBRE). The analysis excludes funding through the IDeA mechanisms, i.e. CoBRE, INBRE and IDeA-CTR grants. For each year, the percentage of NIH research project funding awarded to the institution was calculated. All data was normalized to 1 for the proportion of NIH research project funding at the institution in Year = - 5 of the analysis.

