## Supplementary figures and images for "Efficacy of Centers of Biomedical Research Excellence (CoBRE) Grants to Build Research Capacity in Underrepresented States"

### Supplemental Figure 1

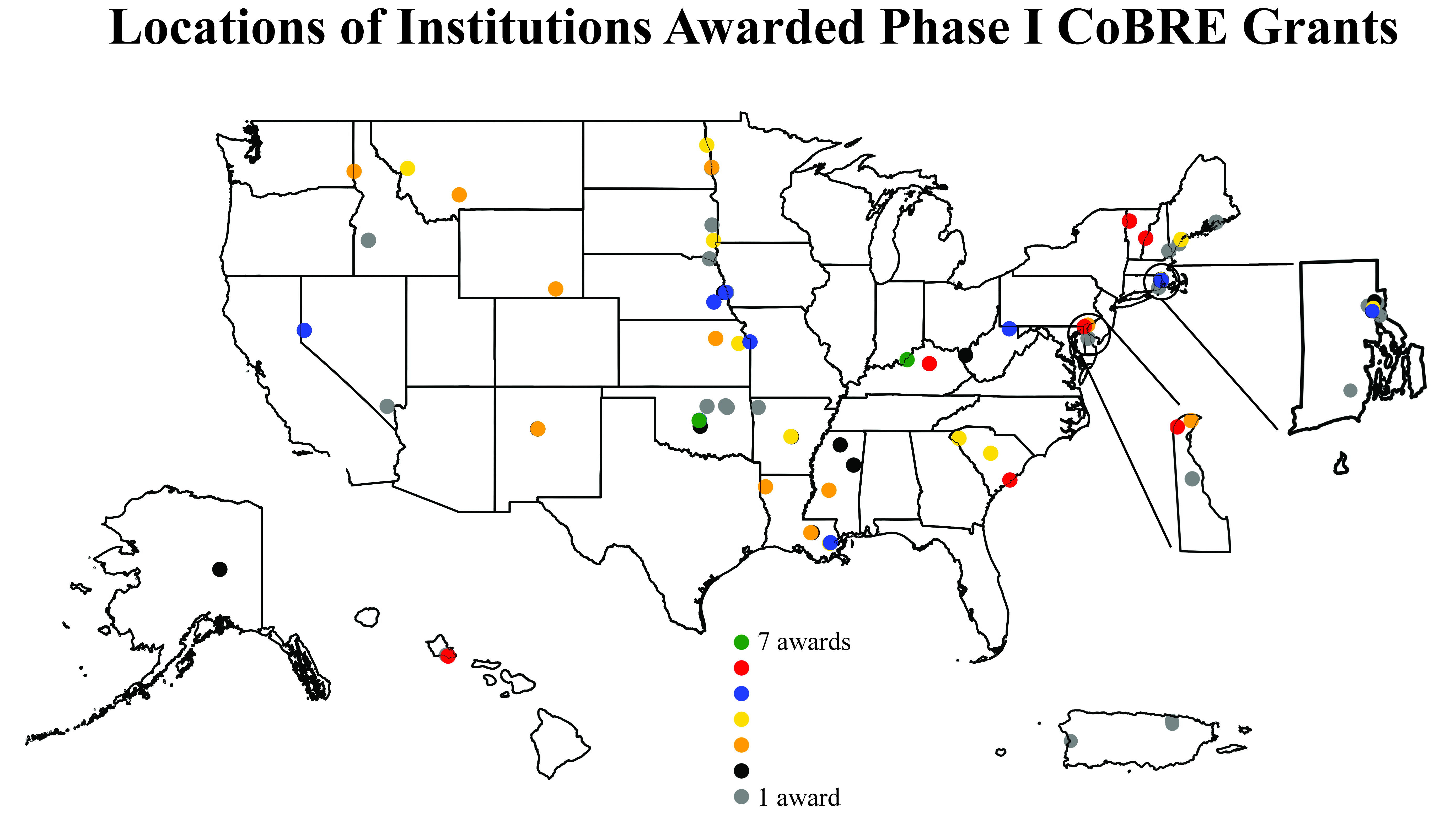

### Supplemental Figure 2

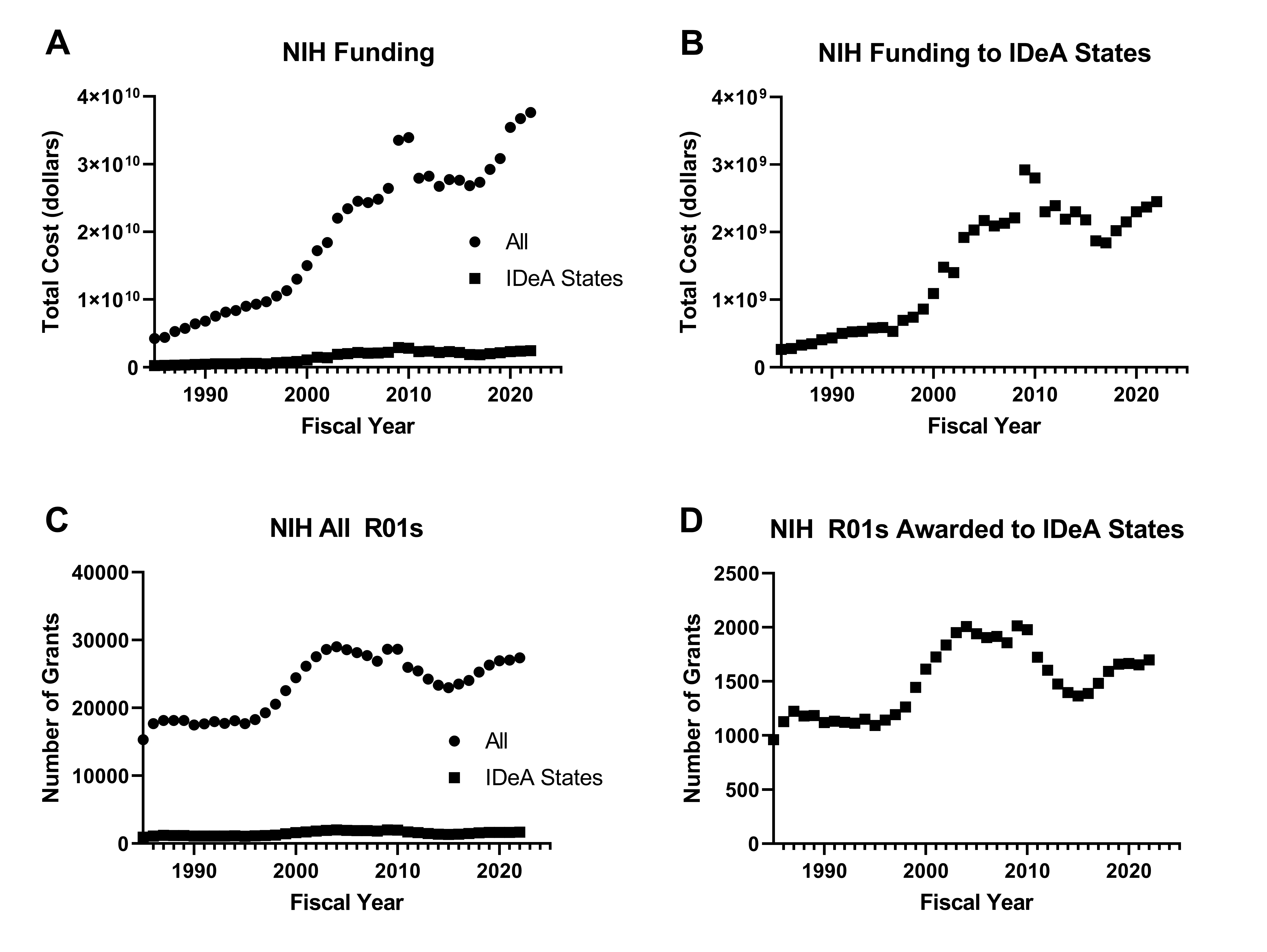

### Supplemental Figure 3A

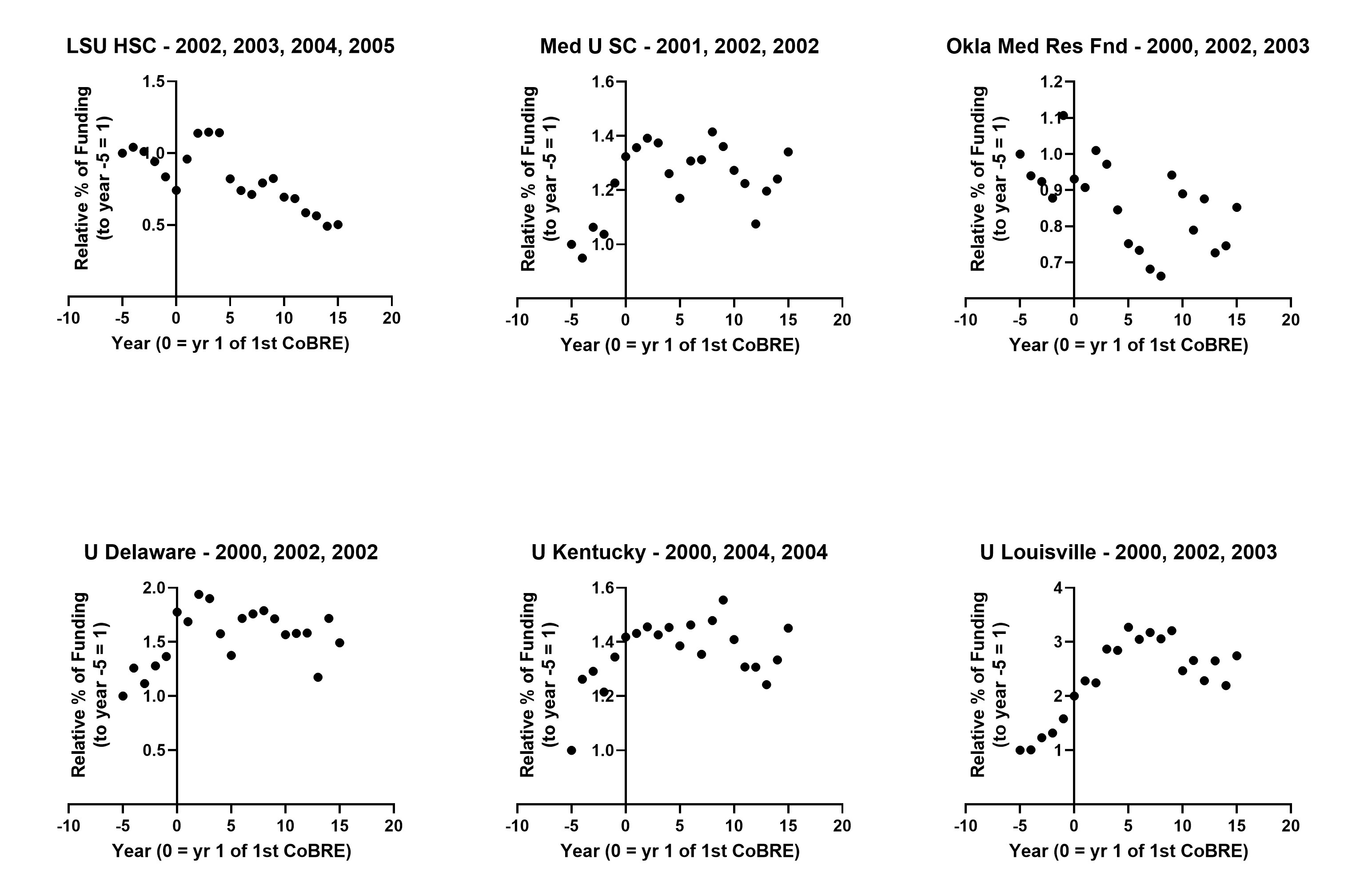

### Supplemental Figure 3B

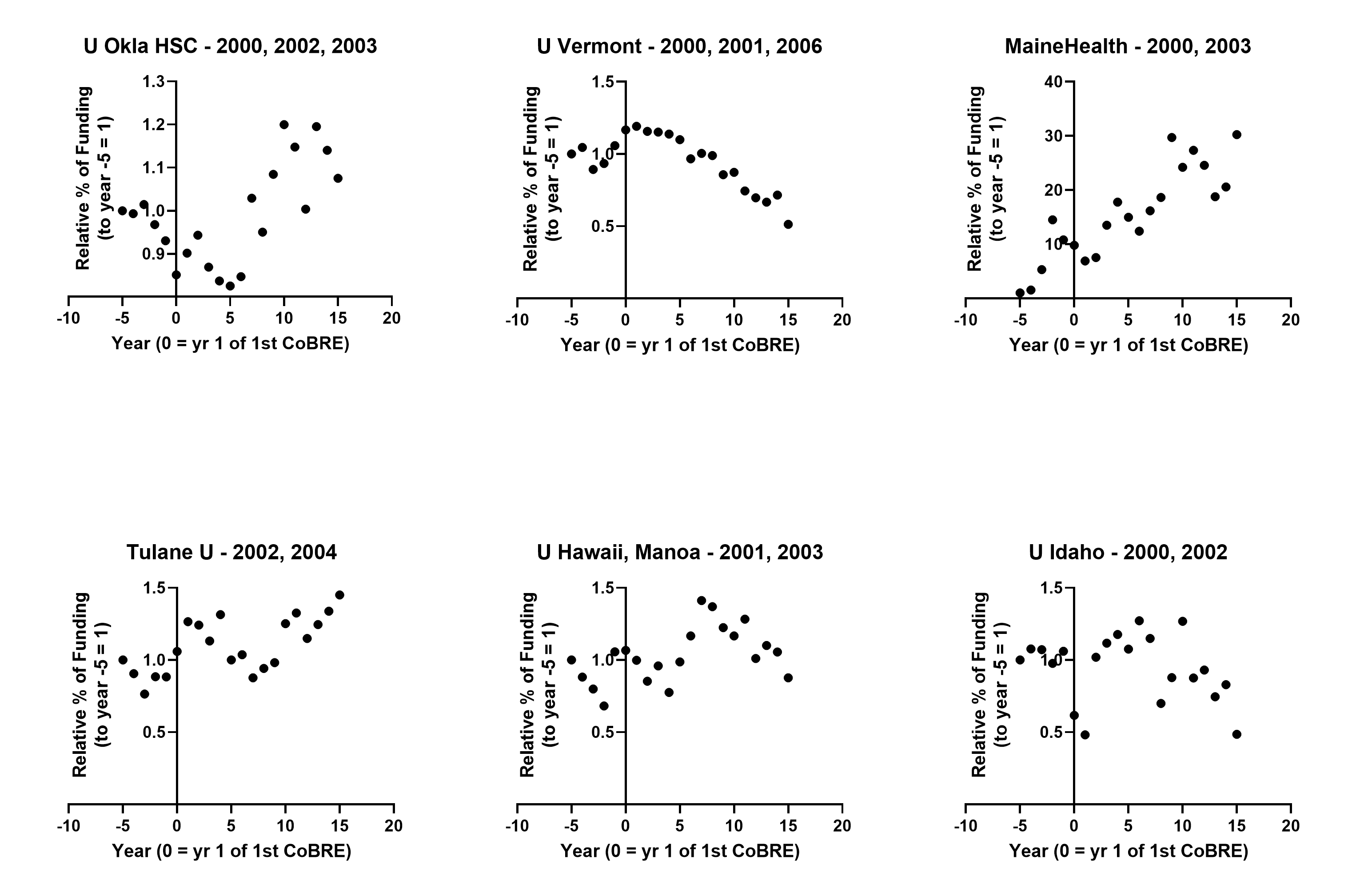

### Supplemental Figure 3C

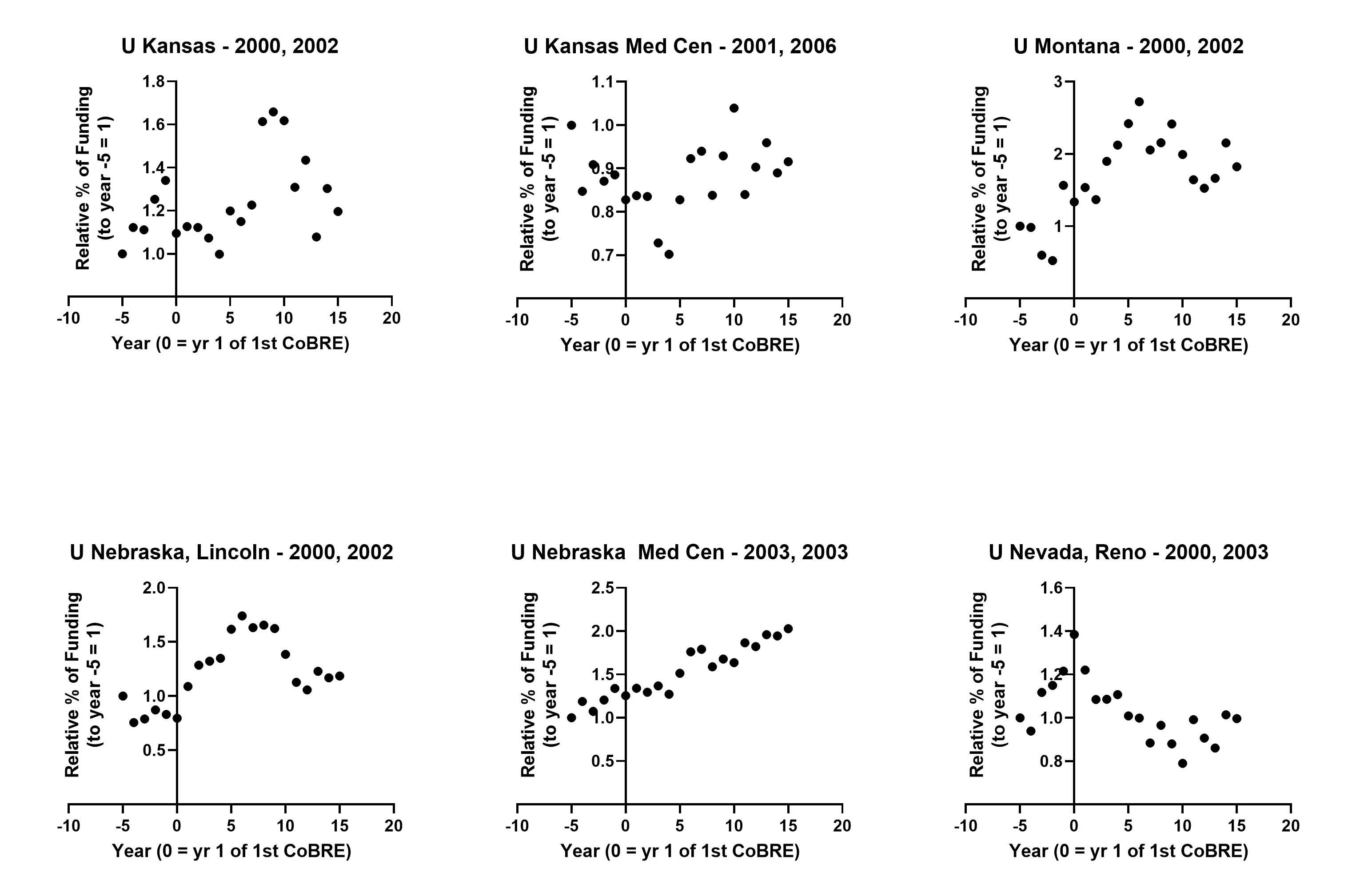

### Supplemental Figure 3D

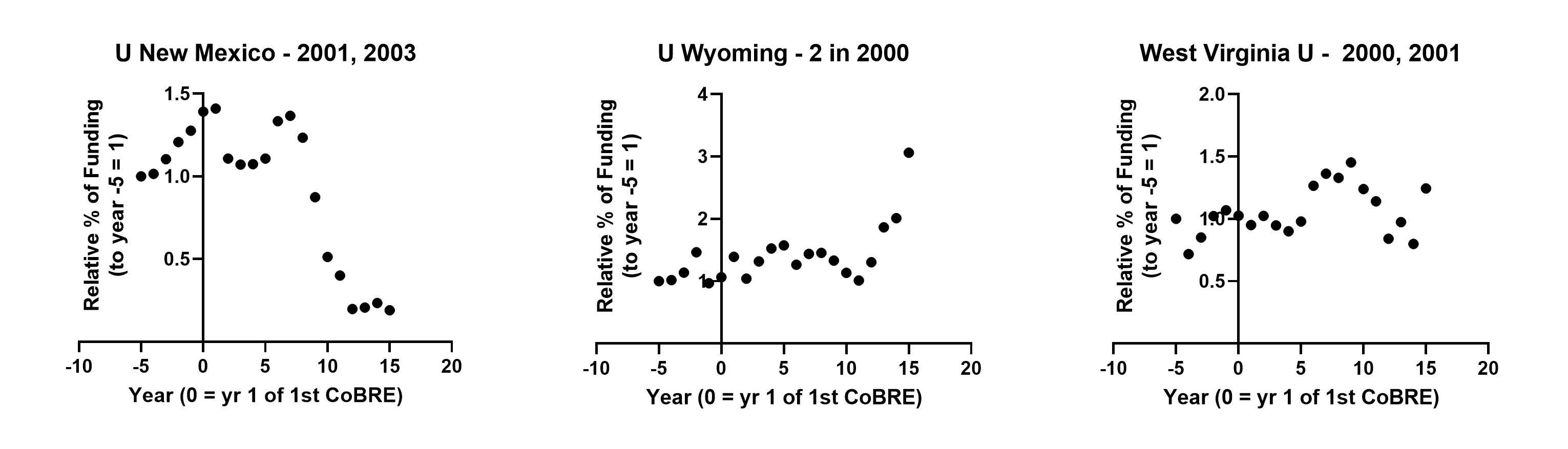
